# Impact of walking speed and motion adaptation on optokinetic nystagmus-like head movements in the blowfly Calliphora

**DOI:** 10.1101/2021.10.18.464799

**Authors:** Kit D. Longden, Anna Schützenberger, Ben J. Hardcastle, Holger G. Krapp

## Abstract

The optokinetic nystagmus is a gaze-stabilizing mechanism reducing motion blur by rapid eye rotations against the direction of visual motion, followed by slower syndirectional eye movements minimizing retinal slip speed. Flies control their gaze through head turns controlled by neck motor neurons receiving input directly, or via descending neurons, from well-characterized directional-selective interneurons sensitive to visual wide-field motion. Locomotion increases the gain and speed sensitivity of these interneurons, while visual motion adaptation in walking animals has the opposite effects. To find out whether flies perform an optokinetic nystagmus, and how it may be affected by locomotion and visual motion adaptation, we recorded head movements of blowflies on a trackball stimulated by progressive and rotational visual motion. Flies flexibly responded to rotational stimuli with optokinetic nystagmus-like head movements, independent of their locomotor state. The temporal frequency tuning of these movements, though matching that of the upstream directional-selective interneurons, was only mildly modulated by walking speed or visual motion adaptation. Our results suggest flies flexibly control their gaze to compensate for rotational wide-field motion by a mechanism similar to an optokinetic nystagmus. Surprisingly, the mechanism is less state-dependent than the response properties of directional-selective interneurons providing input to the neck motor system.

## Introduction

The optokinetic nystagmus (OKN) describes a characteristic pattern of fast and slow eye movements that occur as the entire visual surrounding moves over the retina (Fig. 1A). The eyes saccade against the direction of visual motion and then move slowly with the direction of motion. The OKN complements a range of gaze-stabilization strategies which together have numerous benefits for visual processing, including the reduction of motion blur, lowering retinal slip speed, and facilitating the tracking of stationary objects within a moving scene^1–3^. It is displayed by many species, including both vertebrates (rabbit^4^, pigeon^5^, cat^6^, salamander^7^, flying snake^8^) and invertebrates (mantis shrimp^9^, crab^10^, locust^11^), and has proved a fruitful tool for analyzing the neural circuits of visual motion processing (mice^12,13^, crab^10^, zebrafish^14^, macaque, rabbit, cat, dog, pigeon^15^).

**Figure 1.**
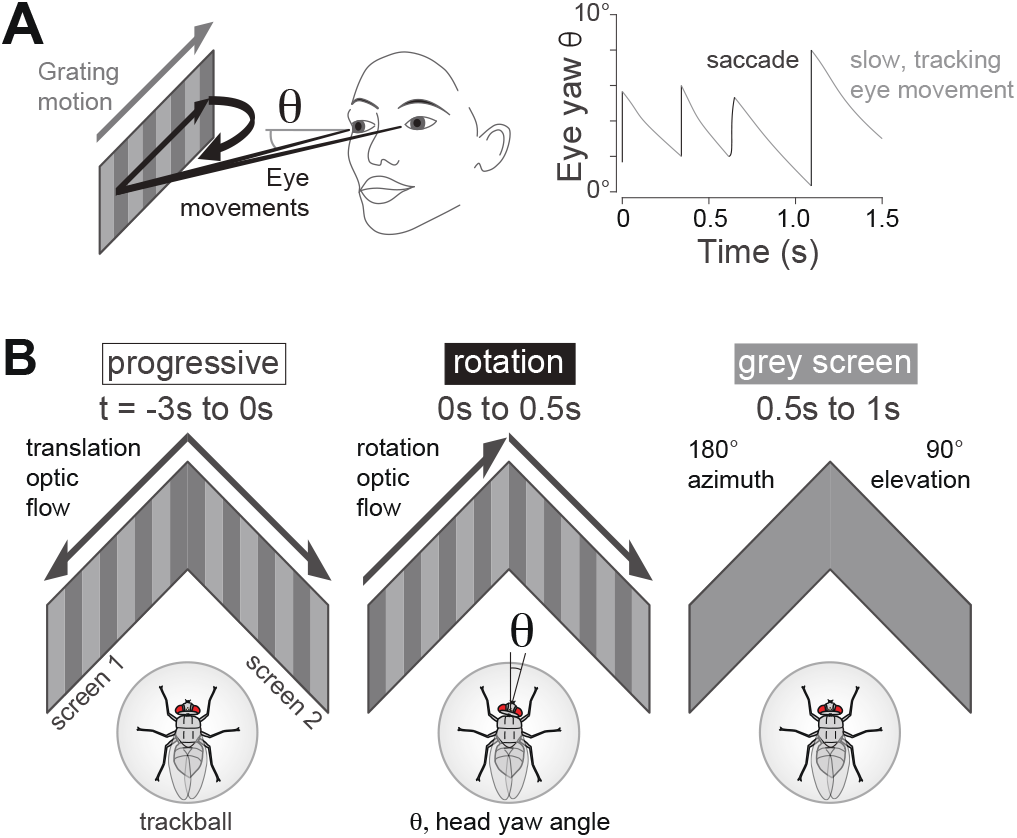
The optokinetic nystagmus and experimental setup and design. **A**. Left: Schematic diagram of human optokinetic nystagmus eye movements in response to a grating moving horizontally left to right (grey arrow). The black arrows indicate the point of fixation moving rightwards as the eyes move syndirectionally with the grating, compensating for its motion. As the yaw angle of the eye’s principal axis (θ) increases towards the limit of its angular range, the eyes saccade leftward, against the motion of the grating. Right: Schematic traces of the yaw eye angle (θ), indicating saccadic increases of the yaw angle against the direction of grating motion, and slow decreases in the yaw angle of the compensatory eye movements that follow the direction of grating motion. The net result is the characteristic sawtooth pattern of yaw eye angle. **B**. Experimental design. Left: In every trial, a fly first viewed a pre-stimulus screen for 3 s (t = −3 s to 0 s) of gratings that moved progressively in a front-to-back direction (‘progressive’) with either one of the following temporal frequencies, 0 Hz, 0.25 Hz, 1 Hz, 4 Hz, 10 Hz, or a grey screen. Center: We measured the yaw angle (θ) of the fly’s head. During the stimulus period (t = 0 to 0.5 s) the fly viewed gratings moving in the same direction simulating the image motion that would occur during a yaw rotation (‘rotation’) with one of the following temporal frequencies, 0.25 Hz, 1 Hz, 4 Hz, 7 Hz, 10 Hz, 13 Hz, 18 Hz, 25 Hz. Right: After the stimulus period (t = 0.5 to 1.0 s), the fly viewed a grey screen. The visual field of the display screens spanned 180° azimuth and 90° elevation.

The neural circuits enabling the OKN can be well studied in flies because much is known about the mechanisms of motion detection at every neural connection between the photoreceptors and the neck muscles in several species. Flies have to move their heads to control their gaze, as their compound eyes are not actuated to perform independent compensatory movements. Large, well-studied interneurons that respond to wide-field visual motion, known as lobula plate tangential cells (LPTCs), innervate directly and indirectly via descending neurons the neck motor neurons that move the fly’s head^16–29^. In particular, LPTCs have motion receptive fields that match combinations of rotational and translational optic flow^30–32^, and are thought to support behavioral responses to wide-field motion, such as head turns and locomotor steering responses^33–36^. These cells integrate motion signals from T4 and T5 cells, which are the first motion-sensitive and directional-selective cells along the pathway^37–40^.

The response properties of the LPTCs are modulated by two aspects related to locomotion. First, locomotion increases their spontaneous activity, response gain and sensitivity to gratings moving with a high temporal frequency^41–45^. The velocity of the visual scene increases with walking speed, and this generates the second modulation of LPTC activity, visual motion adaptation, which decreases the response gain and the sensitivity to high temporal frequencies^46–51^. In moving flies, it is not known how these two aspects balance in LPTCs^52^, and locomotion and visual motion adaptation can affect descending neurons in ways that are different to the effects on upstream interneurons^53,54^. One motivation to characterize the OKN in flies was to identify a visual behavior driven in part by direct connections from LPTCs, whose activity could then provide a read-out for the behavior, especially when the levels of motion adaptation was modulated independently of the walking speed.

Flies rotate their eyes in the horizontal plane with yaw head turns that can be fast and saccadic, or slow^55–60^. While the turning response of the body to wide-field yaw rotations — the optomotor response — has been studied extensively, the vision-induced response of the head has been conventionally studied as a constant head yaw turn rather than the dynamic combination of saccades and slow turns that characterize the OKN^35,61–63^. In two studies in which flying *Drosophila* viewed objects moving against a rotating background, the head position and slow body turns tracked the rotating background while fast, saccadic body turns tracked the position of the object^64,65^. During voluntary body saccades, the head initially turns faster and then slower than the body, thus reducing the time the visual scene moves across the retina^55,58,63^ (but not in walking *Drosophila*^57^). When viewing image motion due to yaw rotation, flies can saccade against the direction of motion^60,61,64^, but it is unclear whether these saccades are part of an OKN to stabilize the visual input, or a casting search for an object to fixate^65^.

Here, we have recorded the head movements of tethered flies standing or walking on a trackball. The flies viewed a period of progressive translational image motion over both eyes that set the level of motion adaptation within a trial, before they were presented with image motion simulating a yaw rotation to induce movements of the head. We find that flies make fast and slow head yaw turns in both directions during the progressive motion stimulus. When they view rotational motion, one the other hand, the fast, saccadic head turns are biased against the direction of motion, while the slow head turns are syndirectional with the direction of the ivisual stimulus, creating head movements similar to an OKN. It is important to note that this response is not a fixed reflex. We observed flies to move their heads only in approximately half the trials, and it does not require the fly to be walking to be engaged, indicating that flies may choose from different gaze-stabilization strategies. In the trials with head movements, the temporal frequency tuning curve follows that found in LPTCs^43,44^, with a peak between 4 and 13 Hz. Surprisingly, we find that the temporal frequency tuning of this OKN-like behavior is robust to changes in walking speed and visual motion adaptation, despite the profound effects these factors have on the temporal frequency tuning of LPTCs.

## Results

To observe the head movement responses to wide-field horizontal motion, we placed tethered blowflies on a trackball in front of gratings that covered a large fraction of the visual field (Fig. 1B) and filmed the head movements with a high-speed camera viewing the fly from above. The yaw rotation stimulus lasted 0.5 s, was preceded by 3 s of horizontal progressive image motion to simulate the horizontal components of forward translation during walking and was followed by a grey screen for 0.5 s (Fig. 1B). As both visual motion adaptation and walking speed modulate the gain and temporal frequency tuning of visual interneurons that encode wide-field motion, we presented the flies with different temporal frequencies of progressive image motion in the pre-stimulus period, randomly ordered selections of 0 Hz, 0.25 Hz, 1 Hz, 4 Hz and 10 Hz, as well as a grey screen with equal mean luminance.

### Head movements in response to rotational image motion

In individual trials, flies moved their head with fast, saccadic turns or with slow turns (Fig. 2). In example trials (Fig. 2A), the saccadic head turns were against the direction of visual motion, shown as a positive change in yaw angle, while slow head turns followed the grating, resulting in a negative change in yaw angle. Combined, these fast and slow head turns have the structure similar to the kinetics of an OKN (cf. Fig. 1A).

**Figure 2.**
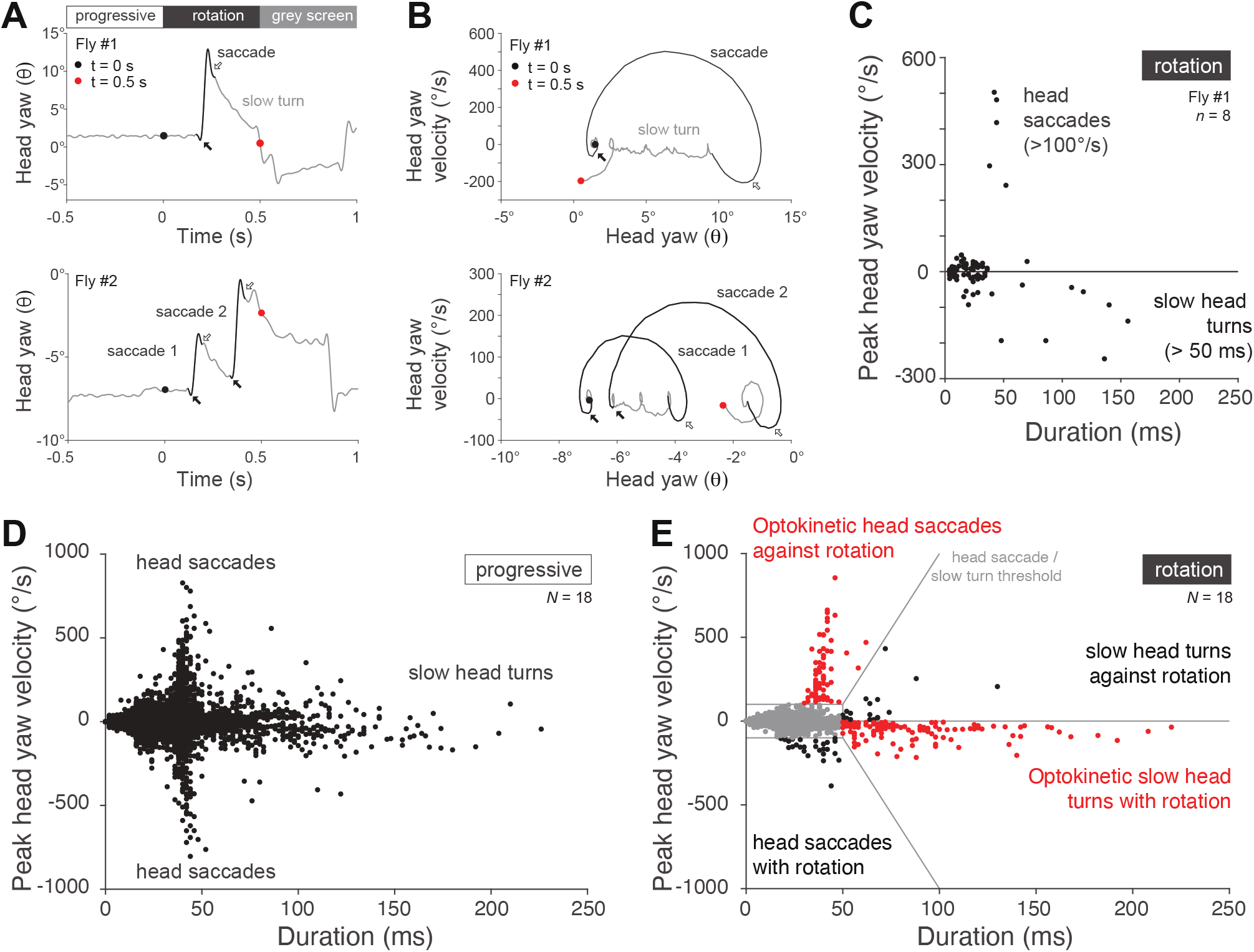
Head movements in response to rotational image motion. **A**. Example trials of head yaw movements of flies viewing 10 Hz progressive motion, followed by 4Hz rotational image motion in the stimulus period. Top: After the onset of the yaw rotation stimulus (black dot), a saccade against the direction of motion (black line) is followed by a slow syndirectional turn (gray line; red dot marks end of stimulus period). Black arrow indicates small counter turn with the direction of motion at saccade start; white arrow indicates overshoot at saccade end where the head again moves with the direction of motion. Bottom: A saccade follows the onset of the yaw rotation stimulus (black line, arrows as before), then a slow syndirectional turn before a second saccade and further slow head turns. **B**. Phase plots showing head yaw velocity and yaw head angle of examples in A. Black and red dots indicate start and end of yaw rotation stimulus period; black and white arrows as in A. **C**. Head turns across all trials for fly #1 shown in A-B for the same pre-stimulus and stimulus conditions shown (*n* = 8 trials). Fast head turns (> 100°/s) are predominantly against the direction of motion. Their duration increases with peak velocity, but last rarely longer than 50 ms. Slow head turns (> 50 ms) are mostly in the direction of the motion stimulus. The magnitude of the peak velocity increases with the duration, but rarely exceeds 200°/s. **D-E**. Peak velocity and duration of head turns of all flies (*N* = 18) for 10 Hz pre-stimulus progressive image motion and 4 Hz stimulus rotational image motion conditions of panels A-C. (D). During pre-stimulus progressive image motion (‘progressive’), saccades and slow head turns occur in both directions. (E). During stimulus rotational image motion (‘rotation’), the distribution of saccades is skewed to positive head yaw velocities and slow turns are dominated by negative peak velocities. Red indicates head turns counting positively towards the OKI, and black indicates those counting negatively towards the OKI. Gray lines indicate thresholds used to classify saccades and slow head turns.

The kinetics of the head movements were revealed by phase plots of the head yaw velocity and yaw angle (Fig. 2B). In this representation, a saccade against the direction of the grating described an upwards and rightwards arc (Fig. 2B, black lines). At the start of the movement, the head turned with the grating for a brief period (Fig. 2B, black arrows) – this was observed as a dip in the head position trace (Fig. 2A, black arrows). The head then accelerated smoothly and peaked in velocity halfway through the turn, before slowing down. In the final phase, the head angle overshot and moved with the grating for a brief period (Fig. 2A–B, white arrows), before the fly engaged in slow turns (Fig. 2B, grey lines). This phase structure of saccadic head turns – the trajectory of the head yaw angle and velocity – was preserved across fast (> 100 °/s) head movements, with peak velocity and duration of the turns covarying for many but not all such turns (compare, e.g., ‘saccade 1’ and ‘saccade 2’ in Fig. 2B; positive fast turns in Fig. 2D–E).

The slow head movements also showed stereotypical structure in the phase plots (Fig. 2B). Sequences of slow compensatory motion were composed of small turns with initial counter-rotations and final overshoot sections of their trajectories, but unlike saccades they frequently lasted longer than 50 ms. The peak velocity and duration of the slow turns (> 50 ms) also covaried, but with peak velocities rarely exceeding 200 °/s (slow head turns in Fig. 2C–E).

To classify head turns, we used thresholds that combined the peak angular velocity and duration (Fig. 2E). These thresholds identified fast, saccadic head turns as having an angular velocity > 100 °/s, and slow head turns as having a duration > 50 ms, separated by linear boundaries. During the pre-stimulus period of progressive visual motion, there were saccadic and slow head movements in both yaw directions (Fig. 2D). During the yaw rotation stimuli, head saccades against the direction of motion predominated, while slow turns were most frequently syndirectional with the motion stimulus (Fig. 2E).

### Rotational image motion selectively recruits OKN-like head movements

The yaw movements of the fly’s head were constrained to a range of ±25° (Fig. 3). The largest head saccades occurred when the head was to the side, with positive saccades occurring at negative yaw angles and vice versa (Fig. 3A Top). Head saccades occurred in both directions for the progressive motion, but the rate of positive saccades - moving against the direction of motion - increased during the rotational motion (Fig. 3A, Bottom – red traces). The rate of positive saccades significantly increased from 0.56 ±0.07 turns/s across all pre-stimulus conditions to 0.79 ±0.10 turns/s for the 4, 7 and 10 Hz rotational motion that generated robust OKN-like responses (mean ±S.E.M.; *p* = 0.004, paired *t*-test, *N* = 18; Fig. 3A, Bottom - red traces). Meanwhile, the rate of negative saccades decreased (from 0.52 ±0.07 to 0.41 ±0.07 turns/s; mean ±S.E.M.; *p* < 0.001, paired *t*-test, *N* = 18; Fig. 3A, Bottom - black traces). Slow head turns also occurred in both directions during the pre-stimulus progressive motion (Fig. 3B, Top, left). Rotational motion decreased the rate of positive slow head turns (0.56 ±0.05 vs 0.30 ±0.03 turns/s; mean ±S.E.M.; *p* < 0.001, paired *t*-test, *N* = 18), and raised the rate of negative slow head turns (0.74 ±0.06 vs 1.25 ±0.11 turns/s; mean ±S.E.M.; *p* < 0.001, paired *t*-test, *N* = 18; Fig. 3B, Bottom). Thus, while fast and slow head movements occur during progressive optic flow, rotational image motion selectively recruits fast head saccades against and slow head turns with the direction of motion.

**Figure 3.**
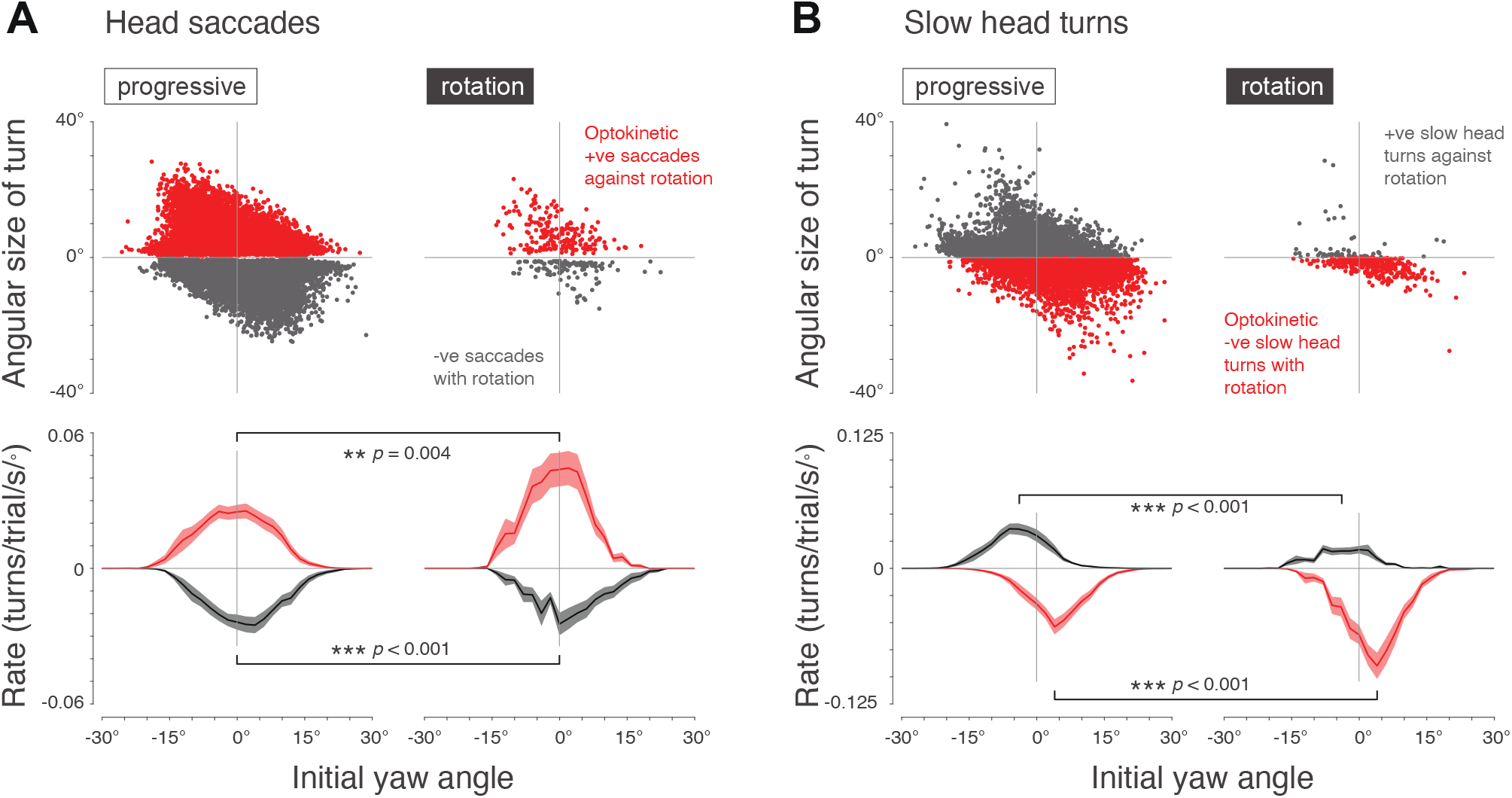
Rotational image motion selectively recruits OKN-like head movements. **A**. Properties of saccadic head turns during progressive image motion of all the pre-stimulus conditions (Left) and during rotational image motion of the 4, 7 and 10 Hz stimulus conditions (Right). Top: the angular size of all positive (red – indicating optokinetic head movement) and negative (black – indicating anti-optokinetic head movement) head saccades, as a function of the initial yaw angle at the start of the turn. *N* = 18 flies. Bottom: the rate of head saccades per trial per second per degree, as a function of the initial yaw angle at the start of the turn. Asterisks and *p*-values indicate significance of statistical comparisons of the rate of saccades per trial per second for all initial angles (paired *t*-tests, *N* = 18; ** *p* < 0.01, *** *p* < 0.001). **B**. Properties of slow head movements, from the same data and using the same plotting conventions as in (A). Here, negative slow movements with the direction of motion are plotted in red – indicating optokinetic head movements – and positive slow movements are plotted in black – indicating anti-optokinetic head movements.

Saccades occurred most frequently when the head aligned with the body axis (yaw = 0°; Fig. 3A, Bottom). In contrast, slow head turns were most often initiated when the head was to the side. For the progressive motion, positive and negative slow head turns most frequently began at yaw angles of −3.5 ±0.9° and 4.6 ±0.7° (mean ±S.E.M.) respectively, values significantly different from zero (positive turns: *p* = 0.001; negative turns: *p* < 0.001, *t*-test, *N* = 18; Fig. 3B, Bottom - left). During rotational motion, the peak initial angles of the slow head turns remained significantly different from zero (positive turns: −3.1 ±1.0°; negative turns: 3.2 ±0.6°, mean ±S.E.M.; positive slow turns: *p* = 0.006; negative slow turns: *p* < 0.001, *t*-test, *N* = 18; Fig. 3B, Bottom - right). Together, these results show that saccadic and slow head turns can be generated at all head yaw angles within the range of movement, and that combinations of fast and slow head movements most often begin with a saccade to the side, followed by a slow turn of the head back to the forward direction.

### Flexible performance of OKN-like head movements, independent of the locomotor state

While the OKN–like responses, consisting of fast turns against followed by slow turns with the direction of motion, occurred for all stimulus conditions, flies did not always engage in this behavior. In some trials, we observed the two kinds of head movements independent of the locomotor state of the animal (Fig. 4A). The distribution of head movements was bimodal (Fig. 4A, Right) and we separated head movements from periods of the head being kept nearly still when the rotation angle was below a threshold value of 2.5°. Likewise, the distribution of walking speeds was bimodal (Fig. 4A, Top). In this case we considered flies to be walking when they exceeded a walking speed of > 3 mm/s, otherwise they were classified as standing (Fig. 4B). In standing flies the head was most frequently still, while walking flies were most likely moving their head (stationary flies, *p* = 0.001; walking flies, *p* = 0.013; Wilcoxon signed rank test with Holm-Bonferroni correction for multiple comparisons, *N* = 18; Fig. 4B). However, we also observed head movements in standing flies, and stationary heads in walking flies (Fig. 4B).

**Figure 4.**
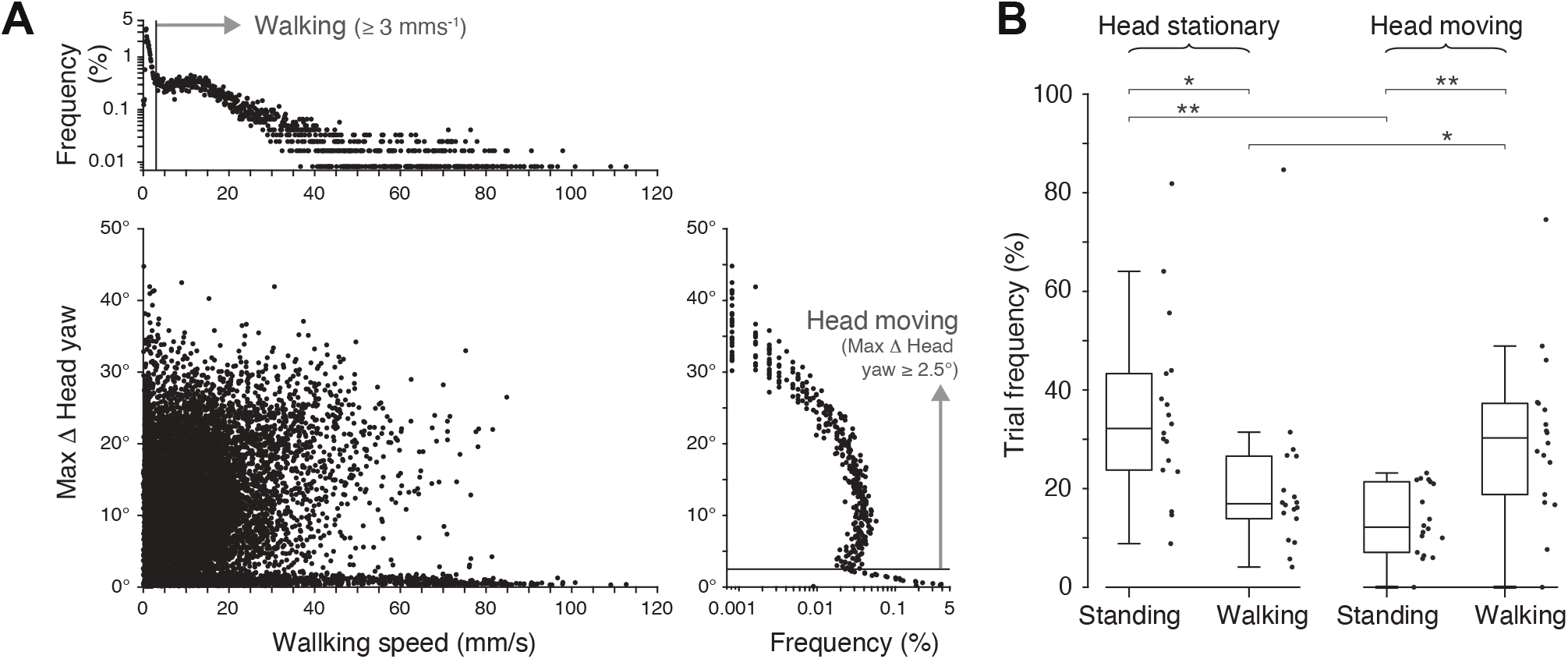
Flexible performance of OKN-like head movements, independent of the locomotor state. **A**. The joint distribution of walking speed and the maximum change of the head yaw angle within an entire trial (4 s), for all flies. Flies were able to move their head or keep them still independently of whether they walked or were stationary. The largest head yaw ranges occurred for the slowest walking speeds, and fast walking flies (> 30 mm/s) either displayed large head movements or kept their heads still. Top: the distribution of walking speeds indicated a bimodal distribution of stationary and walking flies, which we classified by applying a threshold value of 3 mm/s, below which flies were defined to be stationary. Right: the distribution of maximum changes in head yaw angle also indicated a bimodal distribution of flies with heads still or heads moved which we classified by applying a threshold value of 2.5 °/s, below which flies were defined to have stationary heads. **B**. Percentage frequencies of trials in which, from left to right, the flies were: standing with head stationary, 35 ±18%; walking with head stationary, 21 ±18%; standing with head moving, 14 ±7%; walking with head moving, 30 ±17%; all values, mean ±std, *N* = 18. Boxplots indicate the median and quartile ranges, and the whiskers indicate the range of data points that are not outliers. Asterisks indicate statistical significance of comparisons of rates of walking versus being stationary for flies with stationary heads, and moving heads (Wilcoxon signed rank tests, with Holm-Bonferroni correction for 4 comparisons; *N* = 18; ** *p* < 0.01, *** *p* < 0.001).

To quantify the effect of visual motion on yaw head rotations in trials where the fly moved her head, we calculated an optokinetic index (OKI). The OKI is defined as the yaw angle accumulated during fast saccadic head turns minus the accumulated angle due to slow turns during the period of visual stimulation, scaled to a unit time of 1 s (see *Methods*). Thus, if in one second a fly saccades 10° against the yaw grating, and then compensates for the motion of the grating with slow turns for 10°, the optokinetic index is 20°/s. The range of observed head movements was limited to a maximum of ~40° (Fig. 4A, Right). The largest head yaw ranges (max Δ head yaw, Fig. 4A) occurred for the slowest walking speeds, and fast walking flies (> 30 mm/s) either displayed large head movements or kept their heads still.

### Impact of visual motion adaptation on temporal frequency tuning of the optokinetic index, OKI

During the yaw rotation stimulus period, the optokinetic index of walking flies varied with the temporal frequency of the moving grating, reaching a peak between 4 and 10 Hz and decreased at higher temporal frequencies (Fig. 5A). When a grey screen was shown in the pre-stimulus period (Fig. 5A, Left), the response to a yaw rotation stimulus presented at 0.25 Hz contrast frequency is low, the peak response is an OKI of 13.5 ±2.1°/s to a 7 Hz grating which decreases to 3.6 ±1.9°/s for 25 Hz motion. The OKI temporal frequency tuning curves are broadly consistent with of the temporal frequency tuning of visual interneurons responding to wide-field horizontal motion. For example, in response to comparable stimuli the H2 LPTC has a peak response at a temporal frequency of 5 Hz, and a sustained non-zero response to 25 Hz gratings in walking flies^44^. During the pre-stimulus progressive motion period, there was a near-zero optokinetic index, with a small bias between 0.7 and 1.7°/s (mean values for 0 Hz and 10 Hz pre-stimulus conditions, respectively) that was not significantly different from zero except for the 10 Hz pre-stimulus condition (*p* = 0.01, *t*-test with Holm-Bonferroni correction for multiple comparisons, *N* = 18; Fig. 5B).

**Figure 5.**
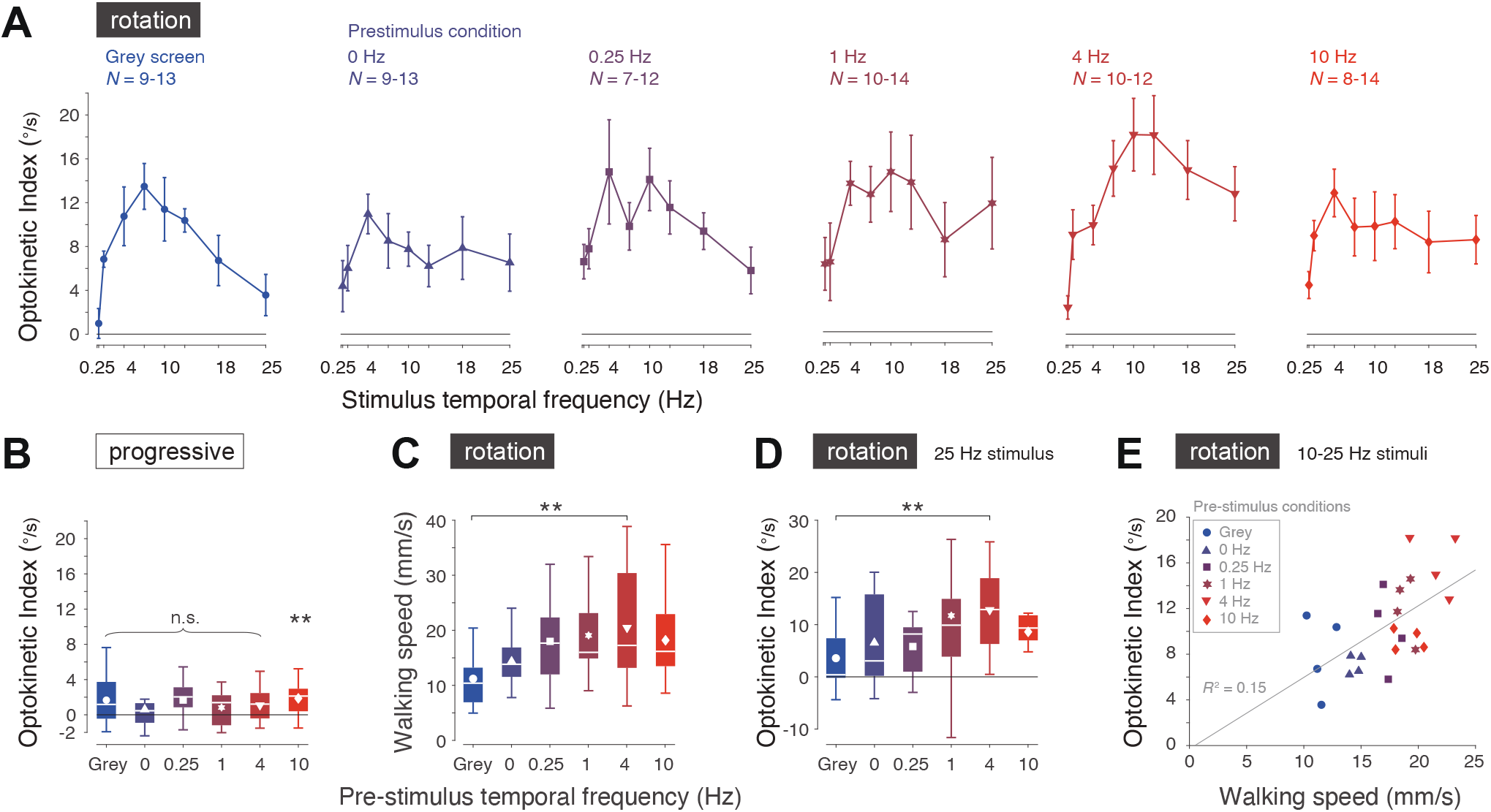
Impact of visual motion adaptation on temporal frequency tuning of the optokinetic index, OKI. **A**. OKI during stimulus period of rotational image motion (‘rotation’) of walking flies (≥ 3 mm/s). The pre-stimulus conditions are labelled in each panel, and indicated by color and plotting symbol. Values were calculated for flies that contributed ≥ 3 trials per stimulus condition. Mean ±S.E.M. shown, with the range of the number of flies contributing trials per stimulus temporal frequency indicated. **B**. OKI during pre-stimulus progressive image motion (‘progressive’) of flies contributing to the data shown in A. Boxplots indicate the median and quartile ranges, whiskers indicate the range of data points that are not outliers, and white symbols indicate mean values. * *p* < 0.05, n.s. not significant, *t*-test with Holm-Bonferroni correction for multiple comparisons, *N* = 18. **C**. Walking speed during the stimulus period of rotational image motion. Plotting conventions as in B. ** *p* < 0.01, paired *t*-test with Holm-Bonferroni correction for multiple comparisons, *N* = 18. **D**. OKI of responses to 25 Hz rotational image motion, for all pre-stimulus conditions. Plotting conventions as in B. ** *p* < 0.01, Wilcoxon signed rank test, *N* = 18. **E**. OKI as a function of the walking speed, for the rotational image motion between 10 and 25 Hz, for all pre-stimulus conditions indicated by color and symbols. (legend). Grey line indicates the linear fit that minimizes the least squared error.

The mean walking speed of the trials varied with the temporal frequency of the pre-stimulus progressive visual motion (Fig. 5C). Flies viewing a grey screen or stationary grating in the pre-stimulus period respectively walked with a forward velocity of 11.2 ±1.3 and 14.4 ±1.2 mm/s during the rotation stimulus (mean ±S.E.M.; Fig. 5C). As the velocity of the pre-stimulus grating increased, the forward walking speed increased significantly to 20.5 ±2.7 mm/s after a 4 Hz pre-stimulus grating (*p* = 0.007, paired *t*-test with Holm-Bonferroni correction for multiple comparisons, *N* = 18; Fig. 5C), and reduced to 18.2 ±1.7 mm/s when the pre-stimulus motion was 10 Hz (mean ±S.E.M.; Fig. 5C). An increase in walking speed is associated with higher sensitivity to visual motion stimuli presented at speeds beyond the LPTC temporal frequency optimum (e.g., > 5 Hz for H2^44^). For high stimulus temporal frequencies, the OKI varied across the pre-stimulus conditions in a way that qualitatively reflected the differences in forward walking speed.

For example, for the 25 Hz stimulus, the OKI increased from 3.6 ±1.9°/s when preceded by the grey screen, to 12.8 ±2.5°/s when preceded by 4 Hz progressive motion (mean ±S.E.M., *p* = 0.009, Wilcoxon signed rank test, *N* = 18; Fig. 5D). This trend of linear scaling of OKI with walking speed was maintained for the responses to 10, 13, 18 and 25 Hz stimuli across all pre-stimulus conditions, with the linear fit explaining 15% of the variance in the OKI (*R*^2^ = 0.15, Fig. 5E).

### Impact of walking speed on temporal frequency tuning of the OKI

To investigate further the effect of walking speed on the OKI, we compared the head movements of flies across all pre-stimulus conditions combined (Fig. 6). During the pre-stimulus progressive motion period, the OKI was near-zero, with mean values between 1.4 and 1.6°/s (compared to zero: 0-3 mm/s, *p* = 0.1; 3-25 mm/s, *p* = 0.002; 25-100 mm/s, *p* = 0.1; *t*-test with Holm-Bonferroni correction for multiple comparisons, *N* = 18; Fig. 6A). In stationary flies (< 3 mm/s), the OKI peaked at 7 Hz, and in slow (3 – 25 mm/s) and fast walking (25 – 100 mm/s) flies the OKI peak occurred at 10 Hz (Fig. 6B). The walking speed also significantly increased the magnitude of the peak OKI, from 15.3 ±2.6°/s in stationary flies to 23.7 ±2.7°/s in fast walking flies (*p* = 0.04, two-sample *t*-test, *N* = 18; Fig. 6B). At high temporal frequencies such as 18 and 25 Hz, the gain of the OKI was not significantly affected (25 Hz: *p* = 0.84; 18 Hz, *p* = 0.84, with Holm-Bonferroni correction for multiple comparisons; two-sample *t*-test, *N* = 18; Fig. 6B). Within the pre-stimulus conditions these trends were maintained, although with less statistical power. Together, these data indicate that the net amplitude of OKN-like head movements is correlated with walking speed, but that walking did not significantly increase the sensitivity to high temporal frequencies.

**Figure 6.**
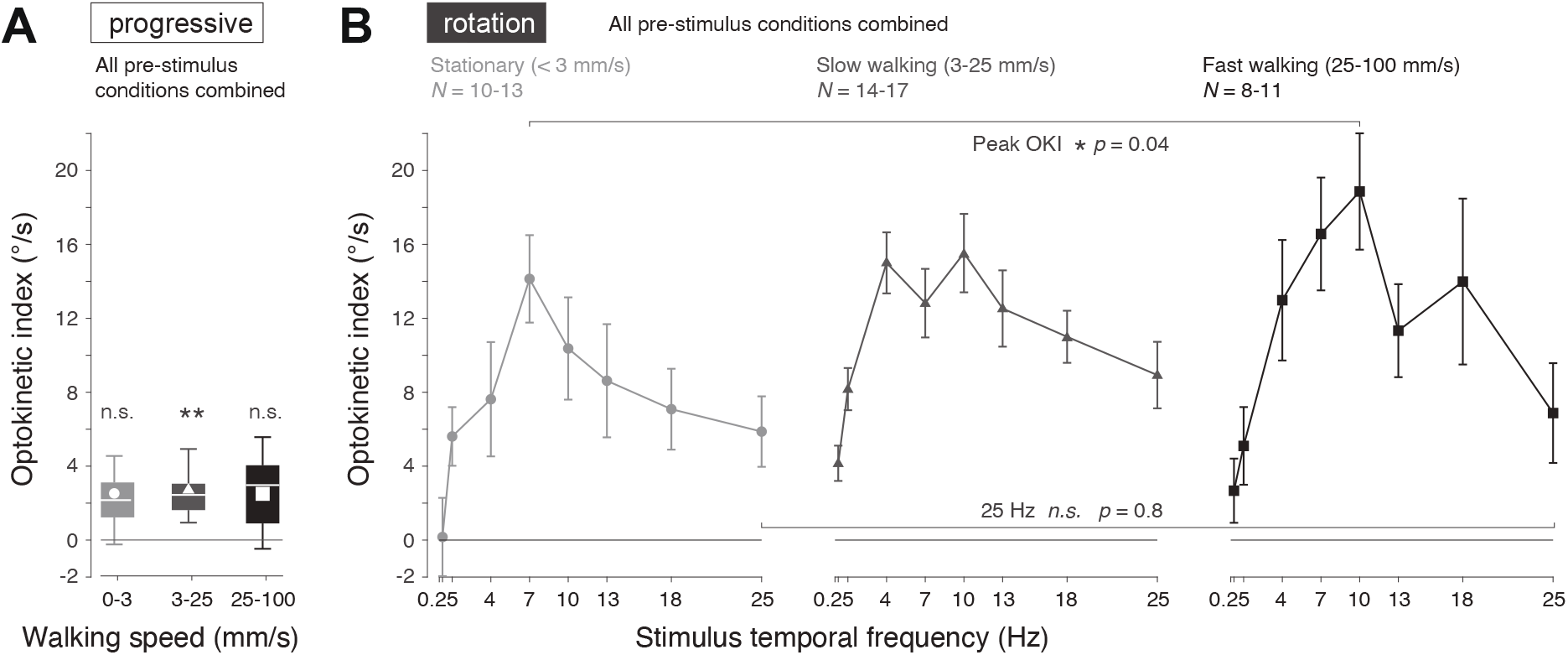
Impact of walking speed on temporal frequency tuning of the OKI. **A**. OKI during pre-stimulus progressive image motion (‘progressive’), for all flies that contributed trials to the OKI during the stimulus period of rotational image motion, shown in B, grouped by walking speeds (left to right): stationary (< 3 mm/s); slow walking (3 – 20 mm/s); fast walking (20 – 100 mm/s). The walking speeds are indicated by the greyscale intensity and symbols. Mean ±S.E.M. shown, and the range of the number of flies contributing trials per stimulus temporal frequency were: *N* = 10-13 (stationary), *N* = 14-17 (slow walking), *N* = 8-11 (fast walking). B. OKI during the stimulus rotational image motion (‘rotation’), for all flies grouped by walking speed as in A. Mean ±S.E.M. shown. The range of the number of flies contributing trials per stimulus temporal frequency were as in A, and indicated. The temporal frequency tuning of the OKI of stationary flies is qualitatively maintained over a range of walking speeds.

## Discussion

We have shown that fast and slow head movements of flies combine to produce OKN-like responses to horizontal image motion, with fast head saccades against and slow head movements in the same direction of the retinal image shift (Fig. 2). In particular, we have shown how the kinematics of saccades and slow head turns can be discriminated in angular velocity and position phase space, allowing small amplitude saccades and slow turns to be detected (Fig. 2B,E). The head saccades were typically generated from central head positions, with slow turns bringing the head back into alignment with the longitudinal body axis (Fig. 3), a sequence of events contrasting with previous characterization of fly head movements as fixed turns in the direction of the rotational image motion^35,61–63^. Head movements were not unconditional reflexes, as flies were able to keep their head still when presented with rotational image motion. The ability to suppress head movements did not require locomotion, in contrast to a previous report on head movement responses to yaw and pitch image motion^62^ (Figs 4B, 6B). The temporal frequency tuning of the combined fast and slow optokinetic head movements matched that of upstream LPTCs reported in other studies^43,44^ (Fig. 5A). While there were modest correlations between walking speed and the magnitude of the OKN-like response (Figs 5C–E, 6), there was little evidence for the profound modulation motion adaptation induces on the temporal frequency tuning or gain of LPTCs.

Our study has focused on yaw rotations, which have specific constraints and functional consequences compared to pitch and roll rotations^16,66^. Previous work has characterized how flying *Drosophila* stabilize their gaze for yaw image motion relative to the wide-field background, even when they produce saccadic body turns to track an object against a moving background^64,65^. Thus, rather than tracking the object by means of gaze shifts, the motion of the object is tracked against a stabilized background. This coordinated head and eye movement strategy has been argued to reduce motion blur and to increase the acceptable input dynamic range of wide-field image motion, allowing the visual system to better track motion in the visual field^56^. Our results are consistent with such a functional interpretation, and we propose that the slow head movements following rotational image motion are part of the blowfly’s gaze stabilization against wide-field background motion.

In *Drosophila*, a recent study has found that saccadic head turns during optokinetic head movements are characteristically preceded by large yaw angles that trigger reset, ballistic head movements against the direction of motion^67^. The kinematics of head movements we observed were stereotypical and consistent with ballistic turns, but in contrast to that study^67^, we rarely observed head saccades from fully rotated head positions (Fig. 3A). This difference may reflect the species studied^55,57^, or the effects on gaze stabilization of flying versus walking^58,59^, but may also be a consequence of the stimuli used or of the task performed. In the vertebrate optokinetic nystagmus, the frequency and amplitude of saccades are affected by the visual stimulus parameters, such as the spatial wavelength of the grating^68^, and by the instructions given to the subject, for example whether to ‘look’ at features in the grating or to ‘stare’ at the grating without trying to follow it^15^. Depending on these factors, saccades can stabilize the gaze with respect to the forward direction^15,69^ or may be triggered by the need to reset the angular position of the eyes to enable the next slow eye movement^15,70^.

In our data, flies were able to perform types of head movements other than those typical of the primate OKN, for example keeping their heads stationary (Fig. 4B) or saccading with the image motion (Figs 2E, 3A). A limitation of our study is that we lack a detailed description of these other types of head movements required to quantify their properties and underlying control mechanisms. For example, Cellini et al. (2021) have suggested that flying *Drosophila* perform oscillatory microsaccades of ~1-2° when viewing stationary visual scenes^67^. These microsaccades, caused by photoreceptor contractions and/or the actuation of eye muscles, suggested to support higher spatial resolution of visual sense^71,72^, may have contributed to many of the small head turns we recorded (Fig. 2D–E), and may be part of a gaze strategy used by the flies when not executing OKN-like head movements (Fig. 4B).

A surprising result of our study indicates that prior motion adaptation does not dramatically reshape the gain or temporal frequency tuning of optokinetic head movements (Fig. 5A), and that the walking speed has only modest effects on the temporal frequency tuning and gain (Fig. 5E, 6A). This finding contrasts with the strong effects of locomotion and motion adaptation on the LPTCs^43,44,46,48,51^. However, it was consistent with recordings of descending neurons sensitive to wide-field visual motion downstream from LPTCs, that also show a lack of visual motion adaptation and a dependence on the animal’s locomotor state^53,54^. The mechanisms underlying such state-independent signaling, however, are still not understood.

Recent work has highlighted how neck motor neurons and LPTCs receive sophisticated signals corresponding to the motor output and anticipated sensory effects of voluntary movements^42,62,63,73–75^. Well-characterized OKN-like head movements offer an opportunity for detailed future studies of the sensorimotor transformations along the comparatively short neural pathways that connect LPTCs and other sensory modalities with the fly neck motor system.

## Methods

### Animal preparation

We used 5 – 16 day old female blowflies, *Calliphora vicina*, from our colony which were kept at 23 °C and at 50% humidity on a 12 hour ON 12 hour OFF light cycle. Flies were briefly cooled and a cardboard tether waxed to the thorax. The wing joints were also waxed to prevent wing movements. A dot of white acrylic paint was applied to the dorsal edge of each compound eye as markers for measuring changes in the head yaw angle. The flies recovered for > 1 hour in a large holding cage with *ad libitum* access to sucrose and water.

We recorded the head movements of 18 flies. Experiments were run at room temperature of 21 °C. Individual flies were used for sets of 64 trials lasting 5 minutes, and then returned to the holding cage while we downloaded movies taken with a high-speed video camera (see below). Flies were tested over a mean of three days, and nine days was the longest period over which a fly was tested.

### Experimental setup

We positioned flies on a trackball system described previously^44^. The polystyrene trackball had a diameter of 48 mm and a mass of the 1.6 g. The mass of the flies was typically ≈ 100 mg, and lighter trackballs were lifted by the flies. The trackball movements were detected by the sensors of two Razer Imperator computer mice (Razer GmbH, Germany), calibrated by rotating a mounted trackball with a step motor along the pitch, yaw and roll axes. The trackball sensors were connected by USB to the controlling computer and driven with NI VISA drivers (National Instruments Corp., USA). The trackball position was sampled at 170 Hz.

The visual stimuli were displayed on two CRT monitors (Vision Master Pro 454, Iiyama, Japan; ‘screen 1’ and ‘screen 2’ in Fig. 1B). The monitors were separated by 80° azimuth, and spanned a visual field of 180° azimuth and 90° elevation (Fig. 1B). Visual patterns were displayed at 170 Hz using PowerStrip (3.90 Entech, Taiwan) and an Quadro NVS290 graphics card (Nvidia Corp., USA), and generated using VisionEgg 1.2.1^74^.

The head position of the flies was filmed with a FASTCAM SA3 high-speed camera (Photron, UK) mounted vertically above the fly. The fly head movements were filmed at 500 Hz with a field of view of 128 × 128 pixels with 8 bits per pixel, controlled by the FASTCAM Viewer software (Photron, UK). Movies were stored on the camera’s internal hard drive and downloaded by an Ethernet connection after each trial.

Experiments were coordinated using Python 2.5 scripts running under Windows XP. The timing of each frame drawn from VisionEgg was used to query the trackball sensors using PyVISA. The filming was triggered at the start of a trial using Python NI-DAQmx package (National Instruments Corp., USA).

### Visual stimuli

Trials lasted for 4 s. For the first 3 s, the flies viewed pre-stimulus visual gratings moving perpendicular to their vertical orientation, in the progressive front-to-back direction simulating forward translation, designed to result in different levels of visual motion adaptation (Fig. 1B). They then viewed for 0.5 s stimulus displays of wide-field horizontal image motion simulating yaw rotations, which were designed to test whether the flies performed head movements similar to an OKN. Finally, they viewed a blank grey screen for 0.5 s, which was included to mitigate the effects of visual motion adaptation between trials. We refer to the first 3 s of each trial as the pre-stimulus progressive image motion period (labelled ‘progressive’ in figures) and the subsequent 0.5 s as the stimulus rotational image motion period (labelled ‘rotation’ in figures).

For all grating stimuli, the contrast (*I*_max_ – *I*_min_ / *I*_max_+*I*_min_) was 50%, and the spatial wavelength was 20° at the closest position to the fly. The grey screen conditions were luminance matched to the mean luminance of the gratings. There were 6 pre-stimulus conditions: a grey screen or gratings moving horizontally and progressively (in the front-to-back direction for the fly) on each screen to simulate the horizontal components of forward translation (Fig. 1B). For the five pre-stimulus conditions with moving gratings, the temporal frequencies were 0 Hz, 0.25 Hz, 1 Hz, 4 Hz and 10 Hz. There were 8 stimulus conditions: gratings moving with temporal frequencies of 0.25 Hz, 1 Hz, 4 Hz, 7 Hz, 10 Hz, 13 Hz, 18 Hz, and 25 Hz.

Trials were organized into sets of 64 trials, in which one fly was shown eight repetitions of all stimulus conditions, for one pre-stimulus condition, in a randomized order. In between the sets of 64 trials, flies were returned to the holding cage.

### Image processing and data analysis

Images were processed using MATLAB 2014a (Mathworks, USA). The angle of the thorax was manually determined for each fly. An intensity threshold was then applied to frames to generate a binary image of the white dots painted on the head, from which the yaw angle of the head relative to the thorax was tracked. The head yaw angle traces, which were sampled at 500 Hz, were filtered with a Savitzky-Golay filter with an order 7 polynomial and a window of 51 data points, corresponding to a 102 ms time window. The head and trackball movement responses to the left-to-right rotation stimuli were inverted and combined with the right-to-left rotation stimuli.

To analyze head turns, we classified every head yaw angle as part of a leftward or rightward turn. By necessity, leftward and rightward head turns must alternate, so we identified all the points where the head yaw direction changed, and classified the head yaw angles between these points as leftward or rightward turns. For each head turn we calculated the duration and maximum velocity of the turn (Fig. 2D–E), the maximum change in yaw angle (Fig. 4A), as well as the dynamics of the turns (Fig. 2A–B). Positive and negative head yaw velocities indicate movements against and with the direction of rotational image motion, respectively. This convention was maintained during the pre-stimulus progressive image motion phase, where, for example, a rightward (clockwise) head turn defines a positive head yaw velocity.

To identify saccades, we used piecewise linear thresholds in a space of head turn duration (x-coordinate) and peak head yaw velocity (y-coordinate) shown in Fig. 2E: for head turn durations ≤ 50 ms, this was a peak head yaw velocity > 100 °/s, or < −100 °/s; and for head turn durations > 50 ms, a point in this space above the line connecting [50 ms, 100 °/s] and [100 ms, 1000 °/s], or below the line connecting [50 ms, −100 °/s] and [100 ms, −1000 °/s]. We classified saccades as turns in one direction, and although saccades are preceded and followed by small turns in the opposite direction, as illustrated in Fig. 2A–B, we did not include these in the duration of the saccade. To identify slow turns, we also used piecewise linear thresholds in the space of head turn durations and peak head yaw velocities shown in Fig. 2E: for turn durations > 50 ms, slow head turns were points in this space between the lines connecting [50 ms, 100 °/s] and [100 ms, 1000 °/s], or below the line connecting [50 ms, −100 °/s] and [100 ms, −1000 °/s].

To identify stimulus periods of trials in which flies moved their heads, we calculated the maximum range of head yaw angles covered, max Δ head yaw, and applied a threshold of 2.5° (Fig. 4A). In trials in which the fly moved her head during the stimulus period, we calculated the optokinetic index by combining the sum of the yaw angle turned, θ, of all the positive saccades, Σθ^+^_*sac*_, and the sum of all the negative slow turns, −Σθ^−^_*slow*_. This combination (Σθ^+^_*sac*_ - Σθ^−^_*slow*_) is the yaw angle covered by OKN-like head movements, that is, saccades against the direction of motion and slow movements with the direction of movement. We then subtracted the optokinetic nystagmus head movements in the opposing direction, the negative saccades, Σθ^−^_*sac*_, and positive slow turns, -Σθ^+^_*slow*_, and scaled this index by the time period assessed, τ:

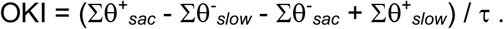

The OKI, therefore, is the angle covered by OKN-like head movements per unit time.

For the data shown in Figs 5 and 6, we used the responses of flies with ≥ 3 trials per stimulus temporal frequency condition.

To identify stimulus periods when the fly was stationary, we calculated the mean walking speed in the stimulus period and also the 0.5 s preceding the stimulus period, to rule out the effect of walking immediately prior to the stimulus period. We used a threshold of 3 mm/s to classify stationary trials (Fig. 4A). During the rotation stimulus, there was no significant optomotor response measured in the rotational velocity of the trackball^77^. This was likely a consequence of the rotation stimulus lasting for only 0.5 s, as comparable yaw rotation stimuli that were longer (4 s) generated robust optomotor responses using the same setup^44^.

### Statistics

To check the normality of data, we used the Anderson-Darling test. For pairwise comparisons of normally distributed data, we used the paired Students *t*-test, and two-sample *t*-test when not every fly contributed data. For testing whether the mean of normally distributed data was significantly different from zero, we used a one sample Students *t*-test. For comparisons between data that was not normally distributed, we used the non-parametric Wilcoxon rank sum test for data where not every fly contributed, and the Wilcoxon signed rank test for pairwise data. To control for multiple comparisons, we used the Holm-Bonferroni correction.

## Acknowledgements

We thank Martina Wicklein for helpful discussions. We thank Graduate School of Systemic Neurosciences at the Ludwig-Maximilian-Universität for administrative support and Axel Borst for internal supervision within the GSN-LMU Master’s degree program for A.S. This work was funded by the Erasmus internship program for A.S. and award FA8655-09-1-3083 to H.G.K from the US Air Force Office of Scientific Research and the European Office of Aerospace Research and Development. K.D.L. was additionally supported by HHMI Janelia Research Campus funding to Michael Reiser.

## Author Contributions

K.D.L. conceived study, contributed to study design, analyzed data, wrote the manuscript, and provided supervision; A.S. contributed to study design, performed experiments and initial data analysis; B.J.H. contributed to study design and provided supervision; H.G.K. conceived study, contributed to study design, critically revised the manuscript, and provided supervision and funding. All authors gave final approval for publication and agree to be held accountable for the work performed therein.

## Additional Information

The authors declare no competing financial and non-financial interests. Correspondence and requests for materials should be sent to K.D.L. or H.G.K.

## Notes

### Competing Interest Statement

The authors have declared no competing interest.

